# Streamlining and Cycle Time Reduction of the Start-up Phase of Clinical Trials

**DOI:** 10.1101/540401

**Authors:** Amani Abu-Shaheen, Ahmad Al Badr, Isamme AlFayyad, Adel Al Qutub, Eissa Ali Faqeih, Mohamad Altannir

## Abstract

**Objective:** The start-up phase of a clinical trial (CT) plays a vital role in execution of novel drug development. Hence, this study aims to identify the factors responsible for delaying the CT start-up phase. Further, it focuses on streamlining and reducing the cycle time of the start-up phase of newly sponsored CTs.

**Methodology:** Thirteen sponsored CTs conducted (between 2016 and 2017) in clinical research department at King Fahad Medical City, Riyadh were analyzed to identify the data specific start-up metrics using **F**ind an improvement area-**O**rganize a team-**C**larify current practices-**U**nderstand source of variation/problem-**S**elect a Strategy-**P**lan-**D**o-**C**heck-**A**ct (FOCUS-PDCA) cycle. Five measures incorporated in the metrics were: date of initial contact with site to the signing of confidentiality agreement; date of receiving questionnaire from sponsor to date of its completion; time taken to review protocol and approve investigational drug service (IDS) form; time taken to review protocol and approve pharmacy and pathology and clinical laboratory medicine (PCLM) form and date of receipt of institutional review board (IRB) submission package to final IRB approval.

Fishbone analysis was used to understand the potential causes of process variation. Mean time was calculated for each metrics prior to and post implementation of the intervention protocol to analyze and compare percentage reduction in the mean cycle time of CTs.

**Results:** Of the various potential factors of delay identified through Fishbone analysis, the two major ones were lack of well-defined timeline for approval and review of the study protocol; and inconsistent IRB meetings. Post introduction of the new intervention protocol, the entire CT lifecycle was reduced by 45.6% (24.8 weeks' vs 13 weeks, before and after intervention, respectively).

**Conclusion:** Varied factors are responsible for delay of the start-up phase of CTs and understanding the impact of each factor allows for optimization and faster execution of the start-up phase of CTs.

## Introduction

Clinical trials (CTs) are essential for testing new drugs, devices and development of new treatment [1]. CTs are widely acknowledged for their role in determining the effectiveness of various therapeutic strategies and diagnostic tests [2]. However, several challenges mask the success of a CT; such as diverse stakeholders (sponsors, investigators, patients, payers, physicians and regulators), infrastructure, logistics, time or other support systems (informatics and manpower) [3]. It has been noted that the start-up phase of a trial sets the tone of the trial and plays an important role in determining the study success. However, starting a clinical trial (CT) is a complicated and time-consuming process. It requires significant understanding of various ethical committees, regulatory bodies and insights into a number of key steps like writing a protocol, applying for funding, and obtaining approvals from all involved stakeholders [4]. Additionally, difficulty in patient recruitment is considered as one of the major reasons of delay in the overall drug development process [2]. The initiation of a new CT usually starts with a research hypothesis, followed by protocol writing, budget and contract negotiation, regulatory essential documents collection, development of a patient recruitment strategy and approvals from the research and development (R&D) department, ethics and other relevant regulatory bodies [4, 5]. On a whole, the start-up phase represents a significant and logistical undertaking [6]. However, duration of the start-up phase generally varies among sites and depends on trial complexity [6, 7]. Lamberti MJ et al. (2013) in their study noted that, early stages of study initiation comprise majority of the lag time where variation in cycle time to the first patient occur by site type (longest for academic institutions and government funded sites and fastest for physician practices) [8]. Krafcik BM et al. (2017) mentioned that 86% of CTs experience delays abiding by the start-up timeline set by the sponsor and contract research organization (CRO) and with a site maintenance cost of up to $2500 per month trial delays can cost the sponsors dearly [6]. From our experience at the research center in King Fahad Medical City (KFMC), Saudi Arabia, the cycle time of start-up phase of any new sponsored CTs is unusually prolonged owing to various factors which negatively impacts the trial conduct. Therefore, the current study aimed to identify the factors that may play a role in delaying the start-up phase of CTs. Also, the study aimed to streamline and reduce the cycle time of the start-up phase of new sponsored CTs.

## Materials and methods

Thirteen sponsored CTs conducted in the clinical research department at King Fahad Medical City, Riyadh, Saudi Arabia in 2016 and 2017 were analyzed to identify the accomplished data specific start-up metrics.

We organized a team of four different disciplines including research center, institutional review board (IRB), investigational drug service (IDS) pharmacy and pathology and clinical laboratory medicine (PCLM) departments. The team was charged with the responsibility to assess the current situation and shorten the time needed for the start-up phase (date of the first contact with the sponsor to the date of getting the IRB approval) of the newly sponsored clinical studies using **F**ind an improvement area-**O**rganize a team-**C**larify current practices-**U**nderstand source of variation/problem-**S**elect a strategy Plan-Do-Check-Act (FOCUS-PDCA) cycle [9]. FOCUS-PDCA is an effective method to systematically solve a simple/complex clinical process problem. It aids in problem solving, change implementation, and continuous improvement in the process [10]. We examined the current process for the start-up phase of new sponsored clinical studies, and used a fishbone analysis to clarify the current knowledge of the process and to understand the causes of process variation.

We used the data available at our site from previous study start-ups to characterize the role of each element of a study in relation to the time required to attain milestones during the start-up phase. The metrics incorporated five measures: (i) the date of initial contact with the site to the date of actual signing of the confidentiality agreement; (ii) the date of receiving the feasibility questionnaire from the sponsor to the date of its completion; (iii) time taken by the IDS pharmacy to review the study protocol and approve the IDS form; (iv) time taken by the PCLM to review the study protocol and approve the PCLM form (v) date of receipt of the IRB submission package by the site through the date of submission to the IRB and date of final IRB approval. The number of days to IRB approval was calculated from time of receiving the IRB submission package from the sponsor to the time between IRB package submission to the final IRB approval.

Mean time for various metrics of clinical trial start-up was calculated before and after implementation of the intervention protocol to analyze and compare the percentage reduction in the mean cycle time for the newly sponsored clinical studies.

## Study interventions

We implemented the following interventions to streamline and reduce the cycle time of the start-up phase of new sponsored CTs: (i) arranging meetings with IRB chairpersons, IDS pharmacy, and PCLM departments to agree on a specified timeline for approving new studies; (ii) completing the IDS form within 7-10 working days; (ii) completing the PCLM form within 5-10 working days; (iv) Modified industry-sponsored research committee for the protection of persons (CPP) approval; (v) assigning lab coordinators to handle all issues related to the send-out lab and get the prices for different tests; (vi) arranging meeting with IRB chairperson, IRB members, and the principle investigator for discussing the protocol and resolving queries prior to the full IRB meeting; (vii) simultaneous submission of study documents to IRB, IDS and PCLM; (vii) express IRB approval was formulized to fast-track the approval process (within 15 days); and (viii) development of standards operating procedure (SOP) for the pathway of new industry-sponsored research.

## Results

The original start-up phase resulted in a serious delay exceeding 24.8 weeks pending IRB submission. This resulted due to deferral in the PCLM and IDS pharmacy approvals (Figure 1). Therefore, in the original start-up phase, approvals from IDS pharmacy and PCLM must be in place before the study package is submitted to IRB. This would help the trial to be on track and prevent any obvious scope of delay for subject recruitment.

**Figure 1:**
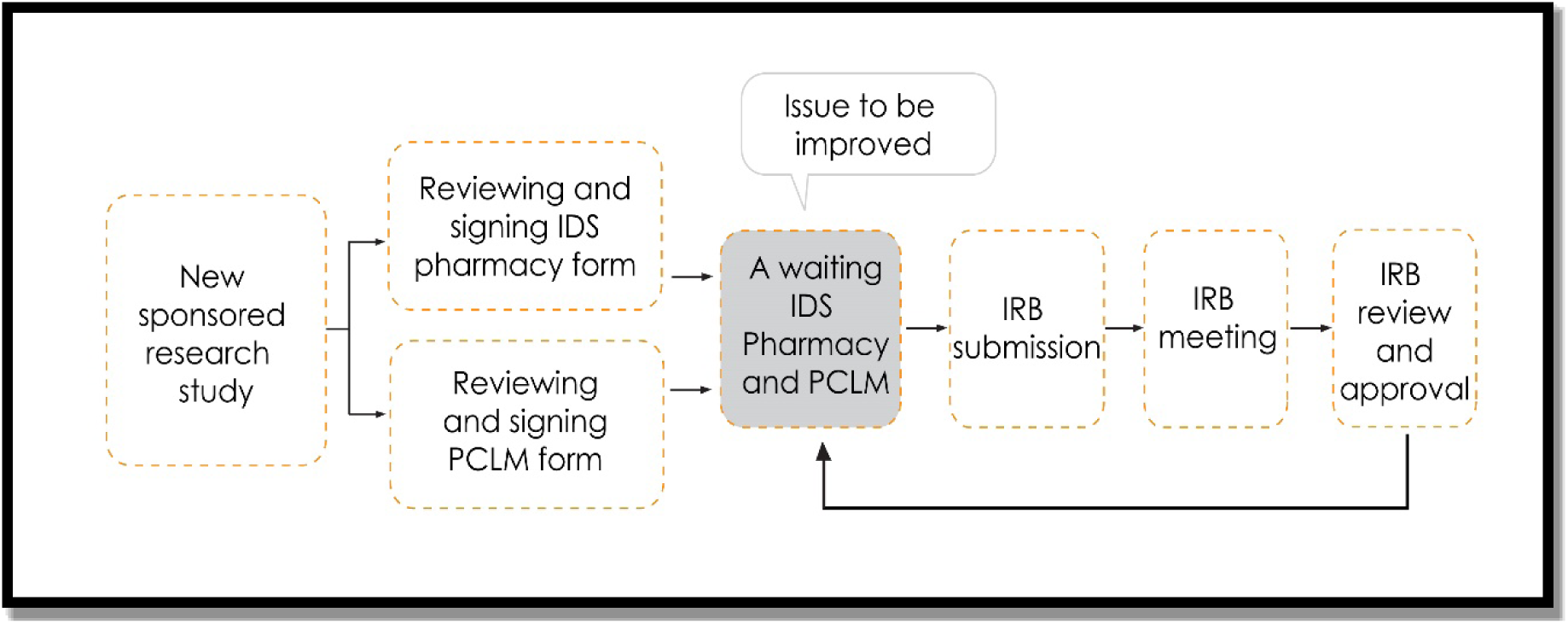
Process-flow of the original start-up phase PCLM: Pharmacy and pathology and clinical laboratory medicine; IDS: Investigational drug service; IRB: Intuitional review board.

Our fish-bone analysis identified numerous reasons for the delay in the current start-up phase and are presented in Figure 2. Two major factors identified were: (i) lack of well-defined timeline for IDS pharmacy, PCLM review, and approval of the study protocol; and (ii) inconsistent convening of the IRB meetings.

**Figure 2:**
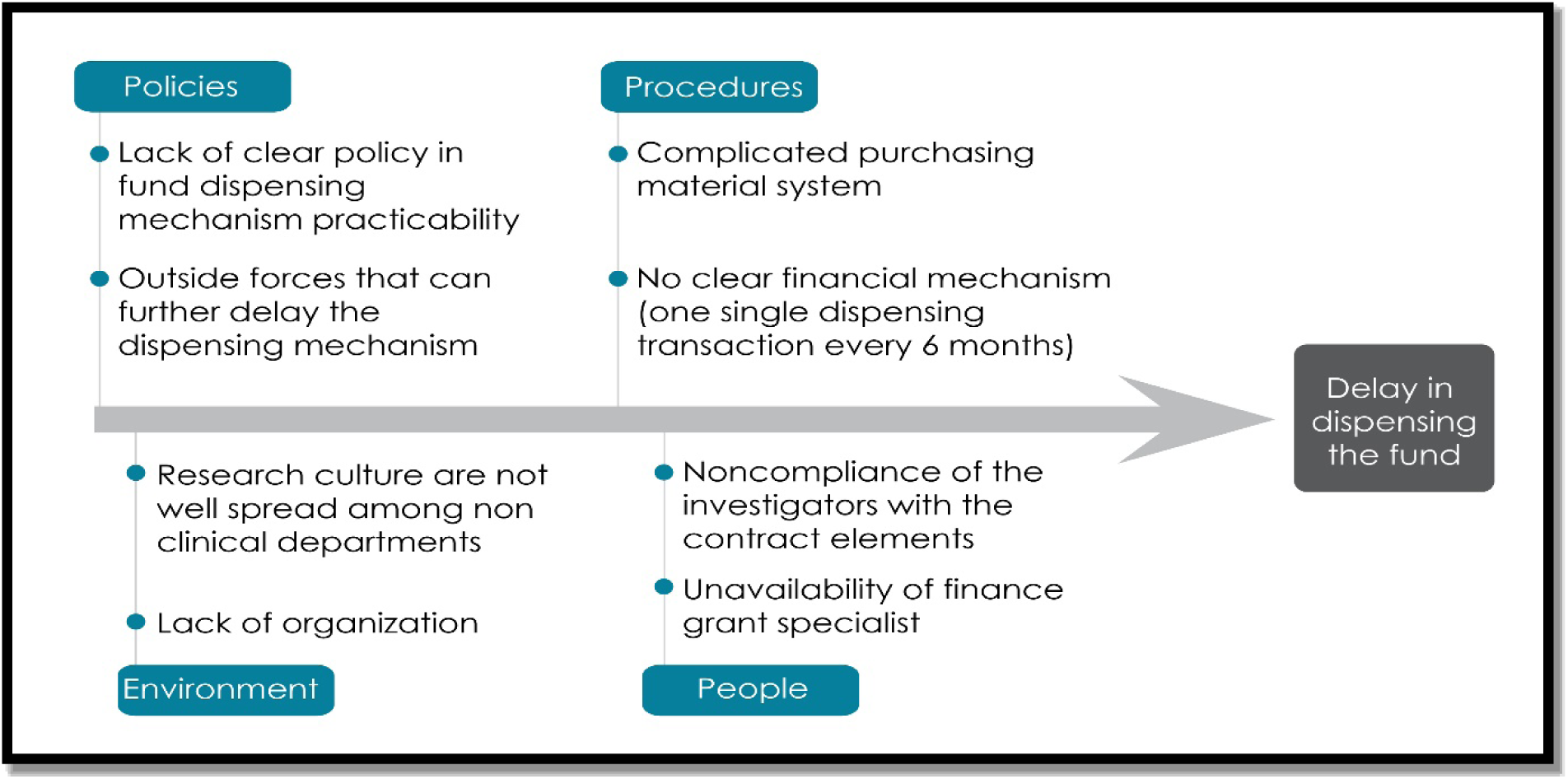
Fishbone analysis of the factors leading to clinical trial start-up delay

Before implementation of the new interventional steps, the mean cycle time of the start-up phase of the new sponsored clinical studies used to be 24.8 weeks. While, after the intervention, the whole process took a maximum of 13 weeks (Table 1). Therefore, with the new interventional protocol, 45.6% shortening of the entire CT lifecycle was noticed (Figure 3).

**Table 1:** Comparison of the mean time for accomplishing different tasks for clinical trial start-up phase before and after implementation of the intervention protocol

|  | **Before implementing**<br>**the intervention**<br>**Mean (week)** | **After implementing**<br>**the intervention**<br>**Mean (week)** | **% of**<br>**reduction** |
| --- | --- | --- | --- |
| **PCLM approval** | 1.7 | 0.7 | 60.4 |
| **IDS approval** | 4.2 | 1.5 | 64.9 |
| **IRB approval** | 7.5 | 6.6 | 12 |
| **From Confidentiality**<br>**Agreement to IRB**<br>**approval** | 24.8 | 13.5 | 45.6 |

PCLM: Pharmacy and pathology and clinical laboratory medicine; IDS: Investigational drug service; IRB: Institutional review board

**Figure 3:**
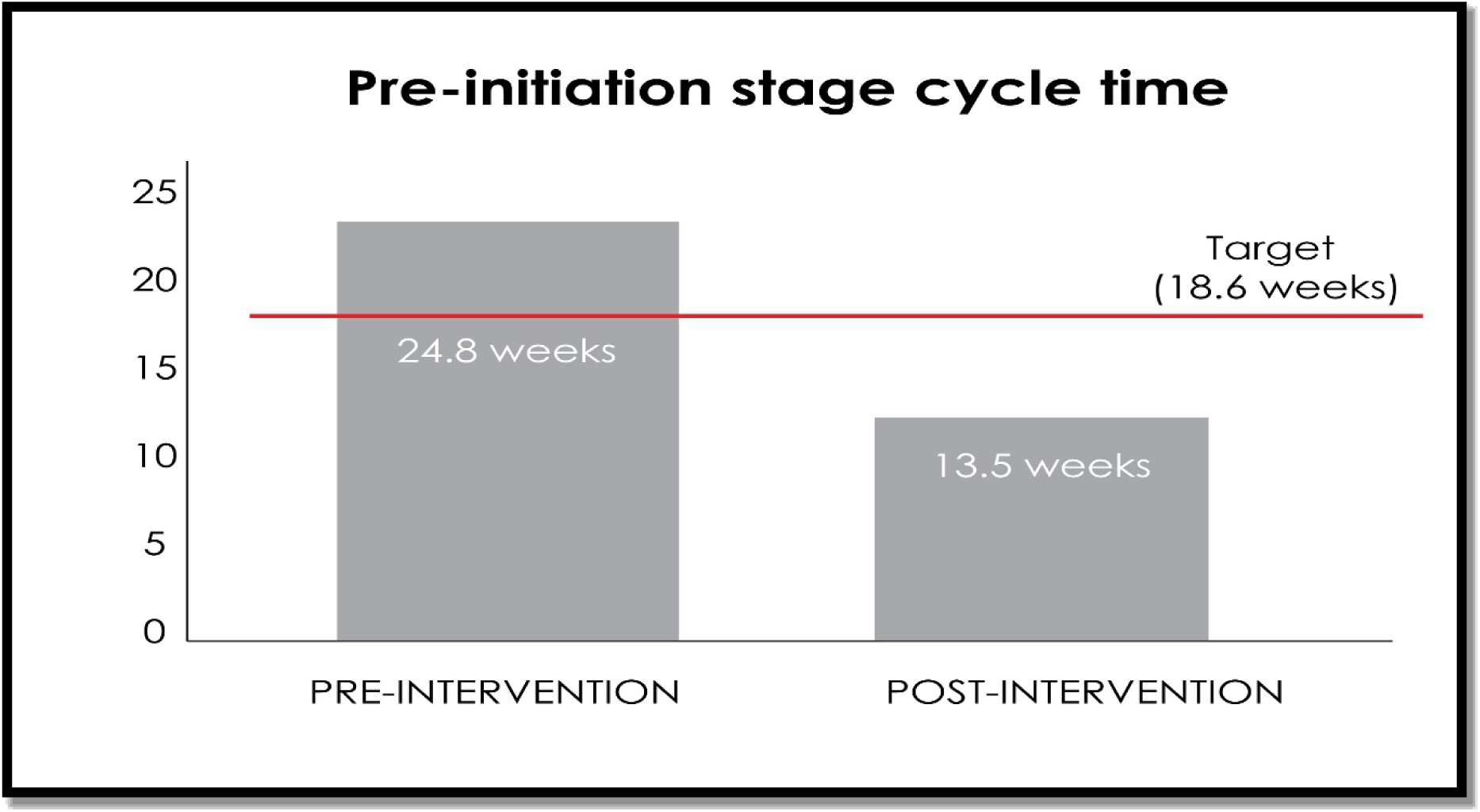
The cycle time of the pre-initiation stage of the new sponsored clinical studies pre- and post-intervention

## Discussion

Findings of our study demonstrated an overall reduction in the mean cycle time of the start-up phase of newly sponsored CTs that gets un-necessarily prolonged due to axillary services required in beginning of the trial. Mainly, inconsistent convening of the IRB meetings, and the extensive period required to obtain approval from IDS pharmacy and PCLM on the study protocol plays as culprit in delaying the start-up phase of a CT. Similar points were also noted by Giffin RB et al. (2010) [3].

While recent studies have reported improved trends in the overall conduct of CTs, sponsors continue to experience significant challenges in meeting overall CT timeline demands [11–14]. Sites must perform several specific activities related to documentation, submissions, agreement approval, and patient visit schedules [4–6]. Historically, the study start-up phase has been viewed as a labor intensive, costly, and time-consuming component of the CT process. Several inefficiencies and limitations continue to threaten the prompt study start-up. Also, impeding efforts in this area is a lack of industry standards regarding the terms or milestones to measure [6].

Krafcik et al reported a faster execution of the start-up phase and better subject enrollment with an earlier IRB approval. [6]. Further, a study conducted by Hurley et al demonstrated that the CT activation period can be reduced through appropriate tools (web-based collaborative workflow tracking tool), staffing, leadership and setting proper priorities. The trail activation time for the six studies used as tests of change were 49, 54, 78, 58, 62, and 32 days. The key activities included during activation phase of the CTs were IRB preparation, Medicare coverage analysis (MCA) processing, Protocol Review and Monitoring Committee (PRMC) review, Medical Research Council (MRC) review, budget negotiation and contract execution. However, delay of more than six weeks observed was mainly due to sponsors [15].

The present study provides a better insight and understanding of the varied steps involved in the start-up phase of a CT. These can efficiently help in reducing the time-lag period in initial CT stages and its associated costs. However, large multi-center studies are required to further support the present findings.

The emergence of newer approaches and strategic intervention protocols to streamline burdensome and time-consuming pre-initiation procedures offer promise, however, with the still uneven adoption of automated and integrated data systems, challenges in predicting start-up timelines and identifying potential holdups will continue. Elimination of the outsider forces to avoid further delay in the pre-initiation stage is a pre-requisite for timely conductance of CT.

## Conclusion

Although notable improvement has been made in the way start-up phase activities are conducted, there remains much work to be done if true efficiencies are to be achieved in CT performance to increase predictability in site start-up.

